# Enhanced *in vivo*-imaging in fish by optimized anaesthesia, fluorescent protein selection and removal of pigmentation

**DOI:** 10.1101/428763

**Authors:** Colin Q. Lischik, Leonie Adelmann, Joachim Wittbrodt

## Abstract

Fish are ideally suited for *in vivo*-imaging due to their transparency at early stages combined with a large genetic toolbox. Key challenges to further advance imaging are fluorophore selection, immobilization of the specimen and approaches to eliminate pigmentation.

We addressed all three and identified the fluorophores and anaesthesia of choice by high throughput time-lapse imaging. Our results indicate that eGFP and mCherry are the best conservative choices for *in vivo*-fluorescence experiments, when availability of well-established antibodies and nanobodies matters. Still, mVenusNB and mGFPmut2 delivered highest absolute fluorescence intensities *in vivo*. Immobilization is of key importance during extended *in vivo* imaging. Here, traditional approaches are outperformed by mRNA injection of α-Bungarotoxin which allows a complete and reversible, transient immobilization. In combination with fully transparent juvenile and adult fish established by the targeted inactivation of both, *oca2* and *pnp4a* via CRISPR/Cas9-mediated gene editing in medaka we could dramatically improve the state-of-the art imaging conditions in post-embryonic fish, now enabling light-sheet microscopy of the growing retina, brain, gills and inner organs in the absence of side effects caused by anaesthetic drugs or pigmentation.

## Introduction

Fish (Zebrafish, *Danio rerio* [1,2]; Medaka, *Oryzias latipes*, [3,4]) have become popular model systems for *in vivo* imaging due to their transparent embryos and their extended genetic toolbox [5,6].

In contrast to the first hours of development, imaging of subsequent stages of development and adult life is obscured by increasing pigmentation and the active movements of the growing animals. Thus, the biggest challenge does not reside on the level of instrumentation, but rather within the specimen itself. They need to be tackled to fully exploit the excellent genetic toolbox. Long term lineaging approaches in both species [7,8] will only deliver dynamic data on fate decisions, once these challenges have been addressed.

This requires a systematic comparative analysis to combine the best suited fluorescent proteins *in vivo* with an improvement of the long-term immobilisation/anaesthesia not interfering with the viability of the organisms and approaches to further enhance the transparency of juvenile and adult fish. All three major challenges are addressed below.

So far, the choice of a given fluorescent protein (FP) as a reporter or tag has often been based on a subjective or pragmatic decision, rather than on quantitative comparison identifying the one best suited. In order to address this, we systematically compared a library of widely used green and red FPs and compared their fluorescence intensities during development upon mRNA injection in medaka and zebrafish embryos.

For this purpose, we scored the respective fluorescence intensities over time and assessed whether a given intensity of a FP is determined exclusively by its amino acid sequence or can be further improved by codon optimization.

The second challenge was identifying a suitable means of anaesthesia for *in vivo* imaging of fish. This is necessary since the standard anaesthetic, Tricaine or MS222, has been shown to only result in incomplete anaesthesia accompanied by adverse effects on heart development (Culver and Dickinson, 2010). Together with Tricaine, we systematically tested the anaesthesia induced by the GABA modulating agent Etomidate and α-Bungarotoxin, a potent nicotinic acetylcholine receptor (nAChR) inhibitor.

The third challenge was overcoming the optical obstructions posed by the pigmentation of eye and peritoneum which prevent the efficient imaging of the underlying cells, tissues and organs. The melanin-containing pigments are not only absorbing light, but are also auto-fluorescent and consequently interfering with fluorescence microscopy [9]. Similarly, the light reflecting properties of the iridescent iridophores interfere with deep *in vivo* imaging. [9]. Both represent a challenge for light microscopy, in particular true for light-sheet microscopy, where the illumination axis is positioned perpendicular to the detection axis [1,2]. Retinal pigmentation completely shields the eye and pigmentation of the head interferes imaging of the brain or any other cranial structure. The most widely used method to suppress the formation of pigment in developing teleost embryos is the application of the toxic and teratogenic drug 1-phenyl 2-thiourea (PTU) [10]. While PTU prevents formation of new melanin pigment, it neither abolishes already present melanin pigmentation, nor blocks formation of iridophore pigment. Consequently, the limited efficacy of PTU towards iridophores together with its severe and toxic side effects highlights the need for alternative experimental approaches.

To overcome these limitations large efforts have been made to systematically breed medaka with reduced pigmentation, suitable for *in vivo* imaging [11,12]. However, since these lines were established by combining several different gamma-ray induced mutant alleles affecting pigmentation, the resulting individuals, mutant in five different loci, are severely affected in life span and fecundity and are consequently challenging to maintain [12]. In order to eliminate pigmentation and light reflection in any genetic background without the need for extensive breeding we devised a CRISPR/Cas9-based knockout strategy to establish pigment-free lines in any background by the parallel targeted inactivation of *oca2* and *pnp4a*. This highly efficient strategy is even applicable for transient approaches upon microinjection.

Taken together, the combination of the approaches presented in this study (double pigment knockout, *α-Bungarotoxin mRNA*, *eGFP* and *H2A-mCherry*) facilitated the imaging of post-embryonic medaka with a MuVi-SPIM (Multiview selective plane illumination microscope) [13,14] highlighting their immediate impact.

## Results

### mVenNB and mCherry are the most intense fluorescent proteins in medaka

In order to assess the fluorescence intensity of different, commonly used FPs (CFP, Clover, eGFP, eGFPvar, eGFPvarA206K, Venus, YFP, mGFPmut2, mVenusNB), we performed a transient ratio-metric approach over extended periods of time. To do so we co-injected equimolar amounts of mRNAs encoding the respective green FP along with the mRNA encoding mCherry as reference. Embryos were injected at the one-cell stage and placed into a 96 well plate for imaging immediately after injection. We determined green and read fluorescence intensities every 20 minutes for at least 40 h at 28°C (exact timeframes are listed in S2 Table). We confirmed that the chorion does not negatively impact on the analysis of injected embryos by comparing embryos with and without chorion at 2, 3 and 4 dpf under the conditions detailed above. This analysis showed no interference of the chorion the determination of the fluorescence properties (Fig 1F, S1 Fig).

**Fig 1.**
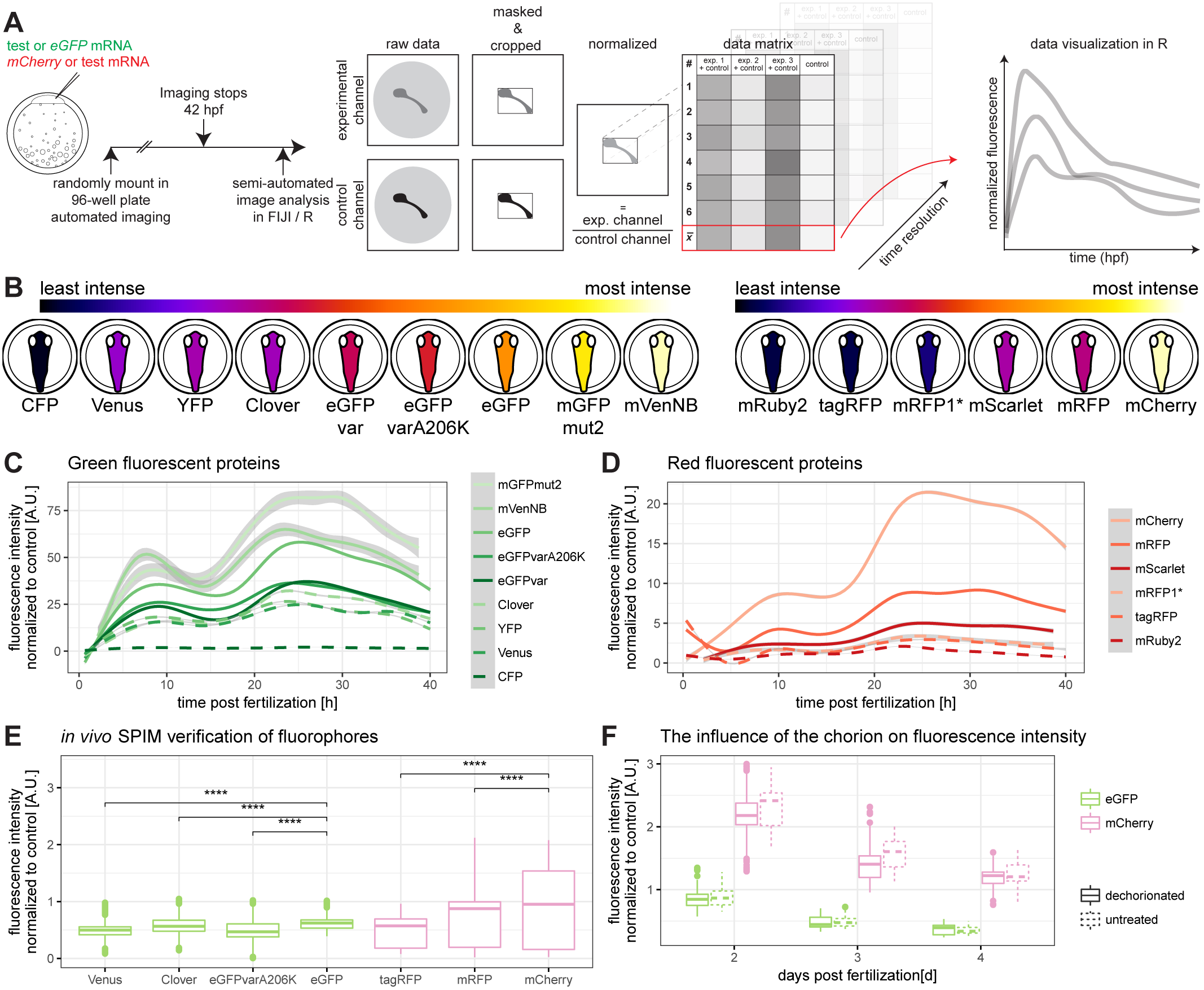
The brightest fluorescent proteins tested in medaka are mVenusNB in the green channel and mCherry in the red channel. (A) Embryos were injected with a green fluorescent protein mRNA to be tested and mCherry mRNA or with eGFP mRNA and a red fluorescent protein mRNA to be tested (1-cell stage modified from [15]). Embryos were mounted in a 96-well plate immediately after injection and automatically imaged for at least 40 h. These data were analysed semi-automated in Fiji and R. Therefore, the raw data were masked and cropped to exclude background. Subsequently, the injection volume was normalized per well by dividing with the value of the control channel at 10 hours post-fertilization. These values were plotted along their biological replicates in R. (B) Schematic overview of fluorescence intensity at 10 hpf, ordered by fluorescence intensity. The lookup table is indicated on the top of the panel, the green fluorescent proteins are shown on the left hand side and the red fluorescent proteins are shown on the right hand side (embryos modified from [15]). (C) Green fluorescent protein fluorescence comparison over time. Plotted is the fluorescence normalized to mCherry per well against the hours post fertilization. Plates were normalized by internal control for fusion of graphs (eGFP). (D) Red fluorescent protein fluorescence comparison over time. Plotted is the fluorescence normalized to eGFP per well against the hours post fertilization. Plates were normalized to internal control for fusion of graphs (mCherry). (E) Verification of previous results by *in vivo*-microscopy of hatched medaka embryos (8dpf) in a light-sheet microscope, comparing the fluorescence intensity of different fluorescent proteins (asterisks indicate P-values: **** P <= 0.0001, *** P <= 0.001, ** P <= 0.01, * P <= 0.05, ns P > 0.05). (F) The influence of the chorion on fluorescence intensity. Exemplary shown are eGFP and mCherry. There is a negligible influence of the chorion of fluorescence intensity measurement. Further information can be found in S1 Fig.

Subsequently, embryos were demounted and image data were analysed in a semi-automated manner by Fiji and R scripts (Fig 1A). In brief we determined the fluorescence intensities by first automatically segmenting the outline of the embryo, followed by the extraction of the mean, median and standard deviation of the resulting region of interest defined by the embryonic outline. The values were normalized to the mean value of the reference mCherry fluorescence intensity at 10 hours post fertilization (hpf). For red fluorescent proteins (mCherry, mRFP, mRuby2, tagRFP, mRFP1*, mScarlet-I), we used eGFP as ratiometric reference and performed an analogous analysis. The resulting fluorescence intensities were ordered allowing instant comparison (Fig 1B). Our systematic analysis revealed that mVenNB and mCherry show the highest fluorescent intensities *in vivo* (Fig 1). We validated particularly high and low scoring candidates by high resolution imaging of mRNA injected embryos at 8dpf in a selective plane illumination microscope (SPIM) (Fig 1E). This analysis confirmed the initial assessments and underscores the impact of autofluorescence of pigment cells.

Previously Balleza and colleagues compared *in vitro* parameters to *in vivo* fluorescence in *E. coli* in order to identify parameters predictive for *in vivo* fluorescence [16]. In order to assess whether this holds also true for vertebrate systems we compared the fluorescence intensities obtained for the different FPs above to their results (Fig 2A). Most FPs show a similar relative fluorescence intensity compared to the normalization value in *E. coli* and *O. latipes*. However, some particularly intense fluorescent proteins such as *mCherry* and *mVenNB* are prominently diverging from the predicted parameters, underpinning the need of an *in vivo* assay to validate novel FPs in the model organism of choice. Similar findings were reported from *C. elegans*, where the *in vitro* parameters are not a sufficient predictor for *in vivo* fluorescence [17].

**Fig 2.**
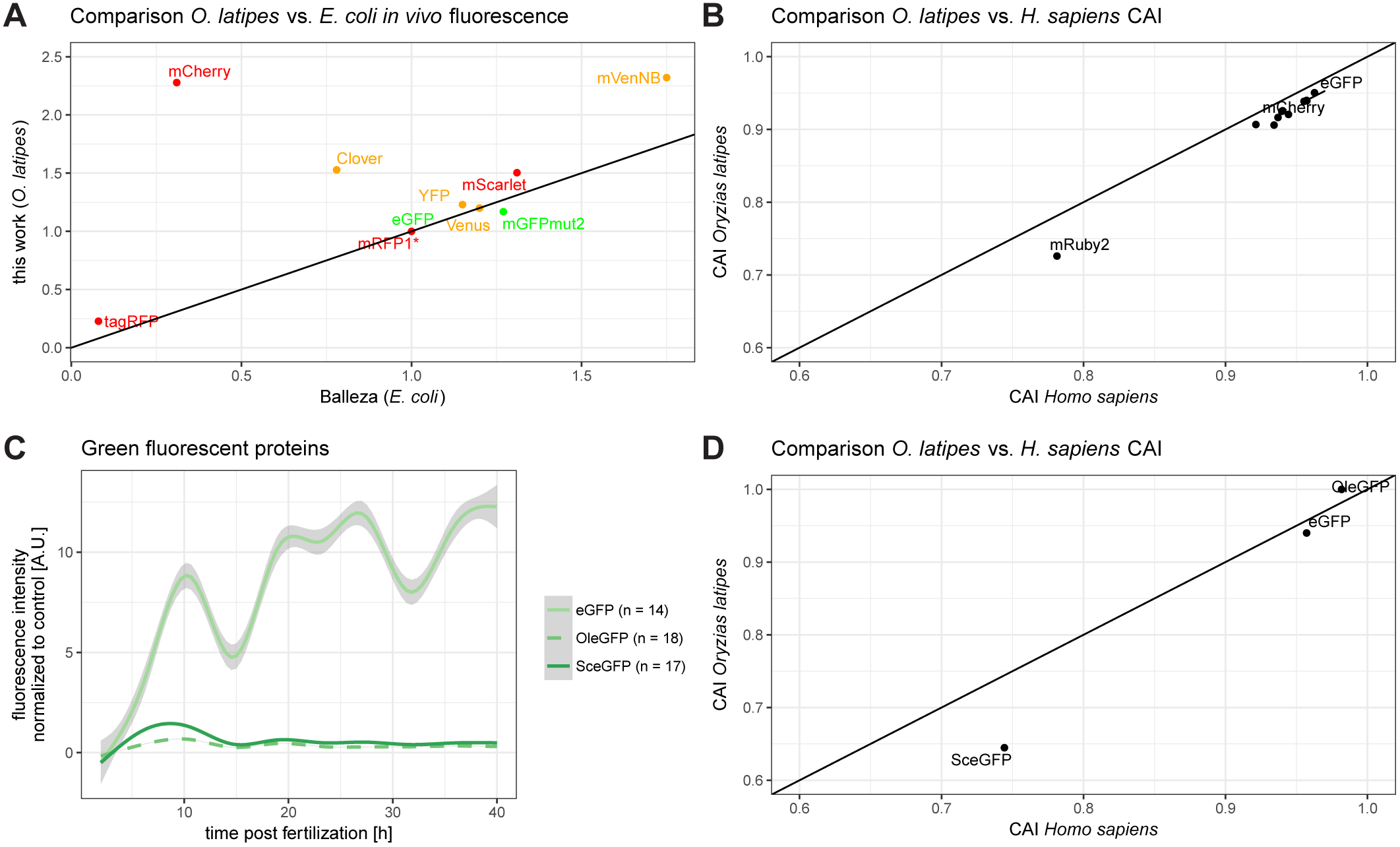
Codon adaptation of *eGFP* for medaka is not improving fluorescence. (A) Relative comparison of *in vivo* fluorescence in medaka with *in vivo* fluorescence in *E. coli*. Indicated are red, green and yellow fluorescent proteins and a diagonal line for orientation. While many of the fluorescent proteins have similar properties, some are significantly different, reinforcing the need of a systematic review of fluorescent proteins in medaka. (B) Codon adaptation indices (CAI) of all prior used sequences. Codon adaptation index of each sequence is plotted for medaka and human along with a diagonal line for orientation. Labelled are the two normalizing fluorescent proteins eGFP and mCherry, and the outlier mRuby2. For a plot with full labelling, refer to S2 Fig. (C) Pure codon usage table driven codon averaging does not improve fluorescence intensity of fluorescent proteins. OleGFP (medaka codon-averaged eGFP) fluorescence intensity is overall lower than SceGFP (yeast codon-averaged) and 25 to 30-fold lower than eGFP. (D) Codon adaptation indices of sequences used in order to score for the effect of codon averaging. Additional to the prior used sequences are an eGFP codon optimized for medaka (OleGFP) and an eGFP codon optimized for yeast (SceGFP).

We addressed a potential impact of protein structure or codon usage on the fluorescence intensities. We first plotted the codon adaptation index (CAI) [18] for medaka and for human for each of the FP sequences used (Fig 2B). To address the question experimentally, we designed codon optimized eGFP for medaka (OleGFP) by OPTIMIZER [19] and plotted it along with an eGFP codon optimized for *Saccharomyces cerevisiae* (SceGFP, Fig 2D). For experimental validation, both mRNAs were injected into medaka embryos, each together with *mCherry* mRNA as injection control and compared to an *eGFP* mRNA control injection. Analysis was performed as described above. The results indicate that codon averaging does not improve the fluorescence intensity. In contrast, the fluorescence intensity of OleGFP is even lower than that of SceGFP, hinting at inefficient translation in case of the codon optimized mRNA (Fig 2C). Taken together, those experiments and the overall similar temporal intensity patterns of all FPs assayed are reflecting a comparable translation efficiency of the different mRNAs and comparable stability of the resulting proteins respectively (Fig 1C-D). Codon optimization does not additionally improve the respective fluorescence properties.

The most intense fluorescent proteins in medaka were also tested in zebrafish and analysed accordingly. Embryos were visualized by the mean over all timepoints, since the trend was not persistent over time (Fig 3A). Green fluorescent proteins (Fig 3B) and red fluorescent proteins (Fig 3C) were normalized to control injections and visualized over time. Strikingly and in contrast to medaka, the trend is not persistent for all FPs, but fluorescence intensity ratios rather vary with time. So, the time of the assay is of crucial importance for the choice of the optimal FP during early zebrafish development.

**Fig 3.**
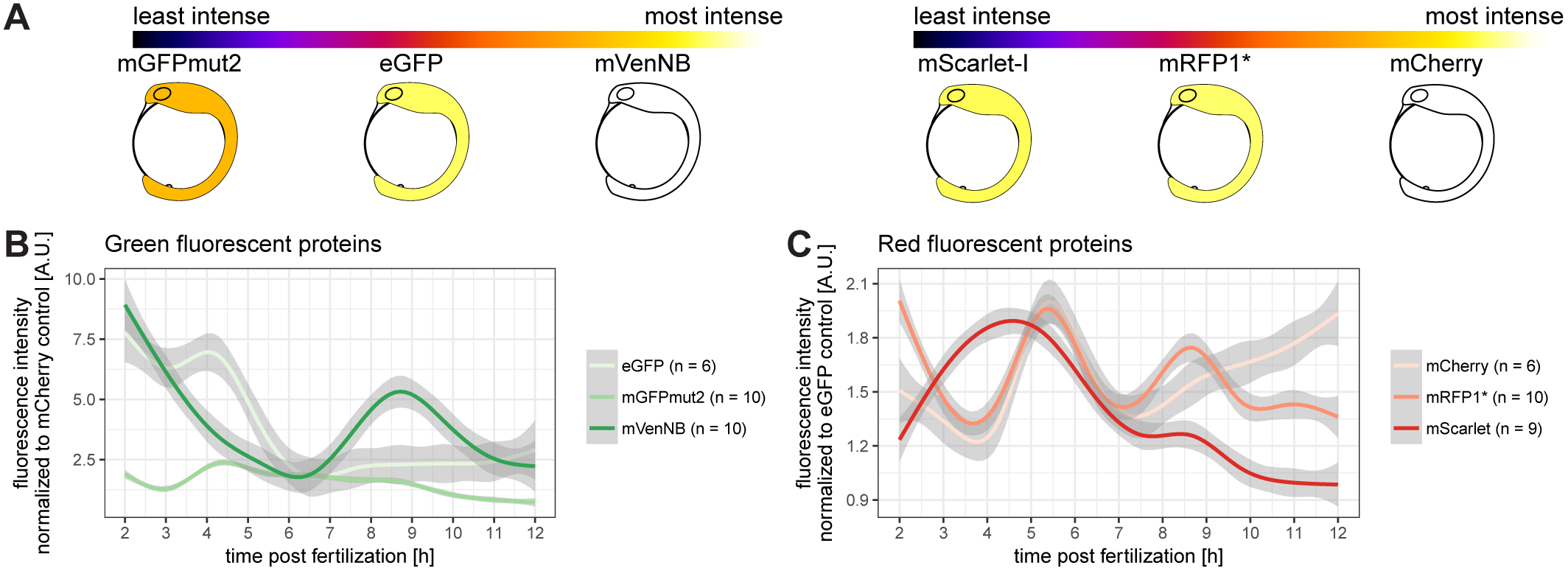
The most suitable fluorescence intensity in zebrafish varies during development. (A) Schematic overview of the mean of the tested fluorescent proteins. The lookup table is indicated at the top of the panel, the green fluorescent proteins are shown on the left hand side and the red fluorescent proteins are shown on the right hand side (embryos modified from [20]). (B) Green fluorescent protein fluorescence comparison over time. Plotted is the fluorescence normalized to mCherry per well against the hours post fertilization. Analysis was conducted as outlined in Fig 1A. (C) Red fluorescent protein fluorescence comparison over time. Plotted is the fluorescence normalized to eGFP per well against the hours post fertilization. Analysis was conducted as outlined in Fig 1A.

### α-Bungarotoxin mRNA injection most efficiently anesthetises medaka embryos

To improve the quality of anaesthesia in medaka, different forms of anaesthesia were compared by scoring movements over time. We either treated embryos with the established anaesthetic Tricaine, or Etomidate or injected one cell stage embryos with mRNA encoding α-Bungarotoxin [21]. All embryos were dechorionated at stage 28 [15], when uninjected embryos were treated with Tricaine, Etomidate, and the respective controls (DMSO, 1xERM). To avoid position effects in the 96-well plate, embryos were loaded randomly and imaged in 20 minute intervals for up to 40 hours. To detect movements, we performed semi-automated image analysis in Fiji [22] and R [23] (Fig 4A). Our analysis workflow delivered a quantitative readout of embryonic movement assessing the efficacy of the anaesthetics (Fig 4B). Strikingly, α-Bungarotoxin was the only anaesthetic efficiently blocking embryonic movement. In contrast, Tricaine and Etomidate did not anesthetize efficiently and treated embryos exhibited strong movements over time (Fig 4B, S1 Movie). Importantly, *α-Bungarotoxin* mediated anaesthetics was reversible and embryos demounted after the analysis eventually showed startle responses, comparable to that of the control treated groups (Fig 4C). S3 Fig gives a visual impression of different projection types of a single well for immediate appreciation. While projection of moving embryos results in a heavy blur, α-Bungarotoxin injected embryos are still visible after projection, with only growth-related movements were detected. Taken together, these results indicate that only *α-Bungarotoxin* functions as effective anaesthetic over extended periods of time. To assess the optimal concentration for reversible anaesthesia suitable for imaging that ensures proper embryonic development, we performed microinjections with different concentrations of *α-Bungarotoxin* (Fig 4D). We determined effective concentrations that are functional even for sensitive lines with limited fecundity facilitating ongoing development and breeding microscopy. In addition we addressed potential effects on cardiac functionality [21], and carefully assayed heart morphology and function after extended imaging in demounted embryos (S4 Fig). Only embryos injected with *α-Bungarotoxin* exhibited normal cardiac function and development, in contrast to the Etomidate or Tricaine treated embryos where heart function was clearly affected by the treatment (S4 Fig).

**Fig 4.**
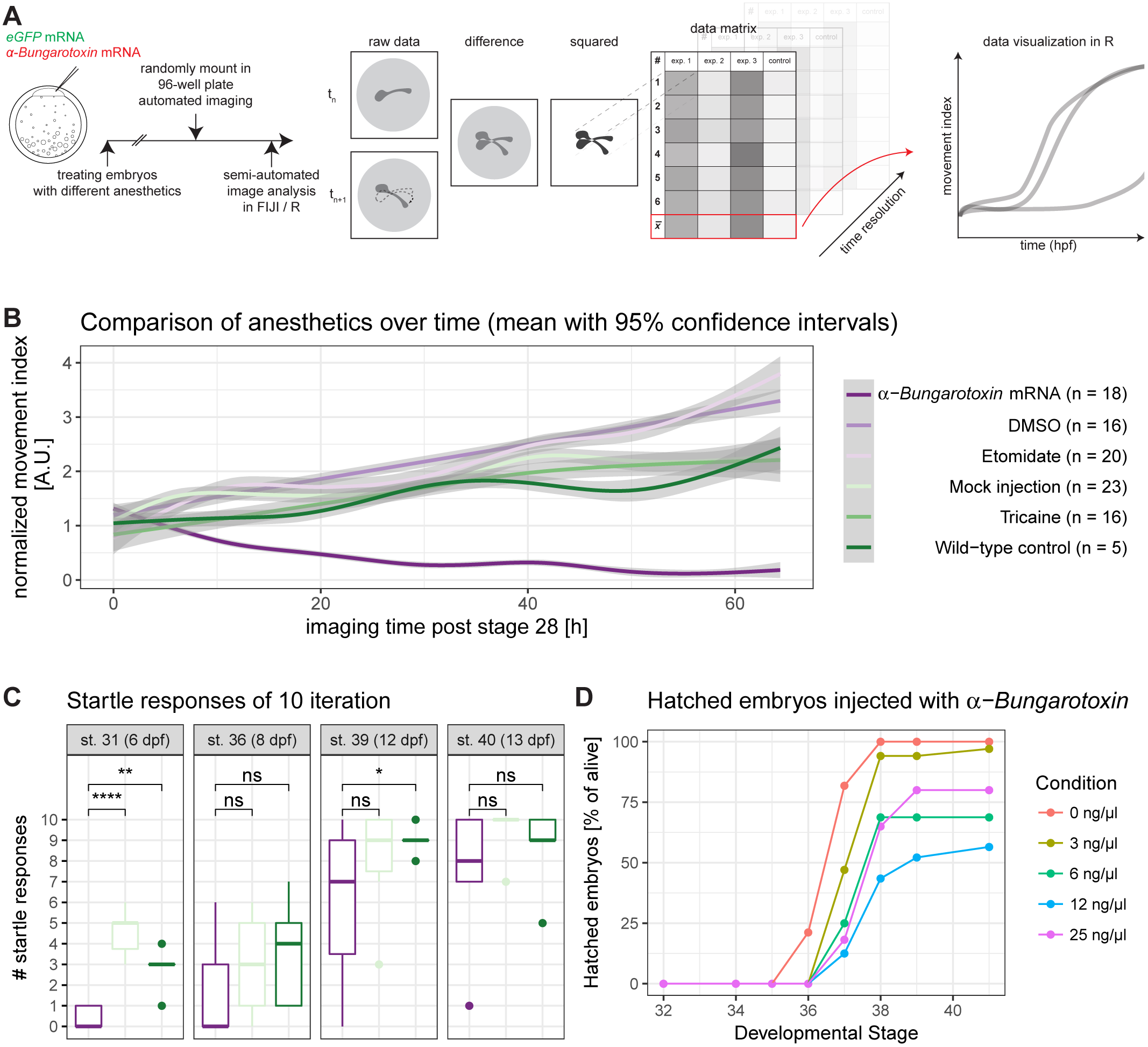
Anaesthesia of medaka is most efficient with α-Bungarotoxin mRNA injection. (A) Embryos were injected with eGFP mRNA or eGFP mRNA along with α-Bungarotoxin mRNA (1-cell stage modified from [15]). Injected embryos and uninjected embryos were dechorionated. Uninjected embryos were treated with Tricaine, Etomidate (experiments), DMSO or 1x ERM (controls). Embryos were randomly loaded in a 96-well plate and imaged automatically. Plates were adjusted by starting stage of imaging and fused into one dataset. Consecutively, image analysis was performed semi-automatically in FIJI and R. Therefore, the difference between each timepoint *n* and its following timepoint *n* + 1 was calculated and squared. These values were imported into R and plotted along with their biological replicates. (B) Movement index of injected/treated embryos over time. Data was analysed as described in A. Strikingly, only α-Bungarotoxin is completely anesthetizing the embryos leading to nearly no detectable difference between the different timepoints (time resolution: 20 min, n as fish per condition is indicated in the legend). (C) α-Bungarotoxin-treated embryos are catching up in startle response ability. Fish were startled 10 times each and the number of responses was noted. Same legend as in Fig 4B applies. (α-Bungarotoxin: n = 12 fish, mock injected: n = 5 fish, wild-type control: n = 8 fish, asterisks indicate P-values: **** P <= 0.0001, *** P <= 0.001, ** P <= 0.01, * P <= 0.05, ns P > 0.05). (D) Different dilutions of α-Bungarotoxin were tested in order to adjust the concentration so fish would wake up after microscopy. Embryos were injected with 0, 3, 6, 12 or 25 ng/μl α-Bungarotoxin mRNA along with eGFP mRNA as an injection control. The number of hatched embryos per condition was scored (0 ng/μl: n = 33 fish, 3 ng/μl: n = 34 fish, 6 ng/μl: n = 16 fish, 12 ng/μl: n = 24 fish, 25 ng/μl: n = 22 fish).

### Efficient CRISPR/Cas9-based double pigment ablation in the injected generation

We employed the CRISPR/Cas9-system in medaka embryos [24] to co-eliminate melanophores in the retina and iridophores two subtypes of pigment cells already in the injected generation. We co-injected of mRNA encoding Cas9 together with sgRNAs targeting a key gene for retinal melanin formation *Oca2* [25] and for the formation of light reflective iridophores, *Pnp4a* [26] to efficiently target melanophores and iridophores simultaneously. Bi-allelic targeting of both loci facilitated by our approach established a high rate (12.5%) of pigment free embryos (Fig 5A) suitable for immediate imaging. The established mutants a fully viable and breed successfully, resulting in translucent embryos and young fish (Fig 5B, S2 Movie). We named the double mutants *spooky*, due to the near-invisibility in petri dishes. These pigment-free fish are amenable to whole mount imaging from embryonic stages up to young adults (S5 Fig). The loss of obstructing pigments in the eye facilitates imaging of the brain by light-sheet microscopy. We combined all approaches described above, injected mRNA encoding the most efficient fluorophores (*eGFP* and *H2A-mCherry* mRNA) into *spooky* mutant fish (*Oca2^-/-^*, *Pnp4a^-/-^*) anesthetized with α-Bungarotoxin and performed long-term imaging of retinal and brain structures previously obstructed by pigmentation (Fig 6). The results obtained underscore the importance of combining all approaches related to the specimen to move whole mount *in vivo* imaging to new frontiers.

**Fig 5.**
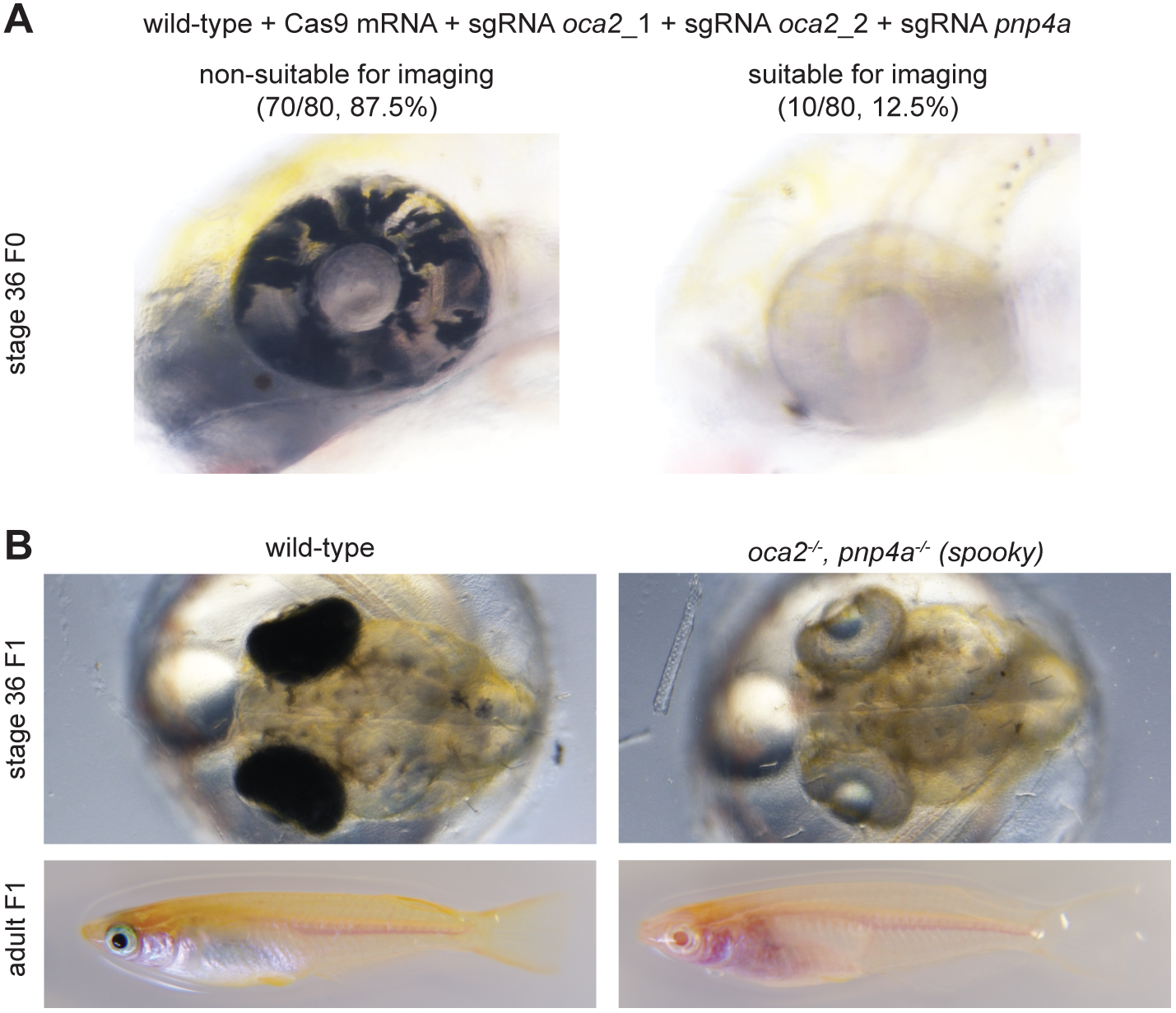
The acute double pigment knockout utilizing the CRISPR/Cas9-system can be deployed in the injected generation. (A) Medaka zygotes were microinjected with Cas9 mRNA, sgRNA (Oca2_1), sgRNA (Oca2_2) and sgRNA (Pnp4a). The ratio of the embryos suitable for imaging was determined (representative examples shown). At least 12.5 % of the injected embryos were suitable for imaging in the injected generation. (B) Injected medaka as shown in panel A were raised, screened and imaged in F1 to compare the effects of sgRNA injection. Wild-type embryos exhibit a dense, black pigment in the eyes, whereas adults additionally exhibit a non-translucent peritoneum. Fish injected with sgRNAs against *oca2* and *pnp4a* show dramatic loss of pigmentation surrounding the eye and the peritoneum, rendering the fish more accessible for microscopy.

**Fig 6.**
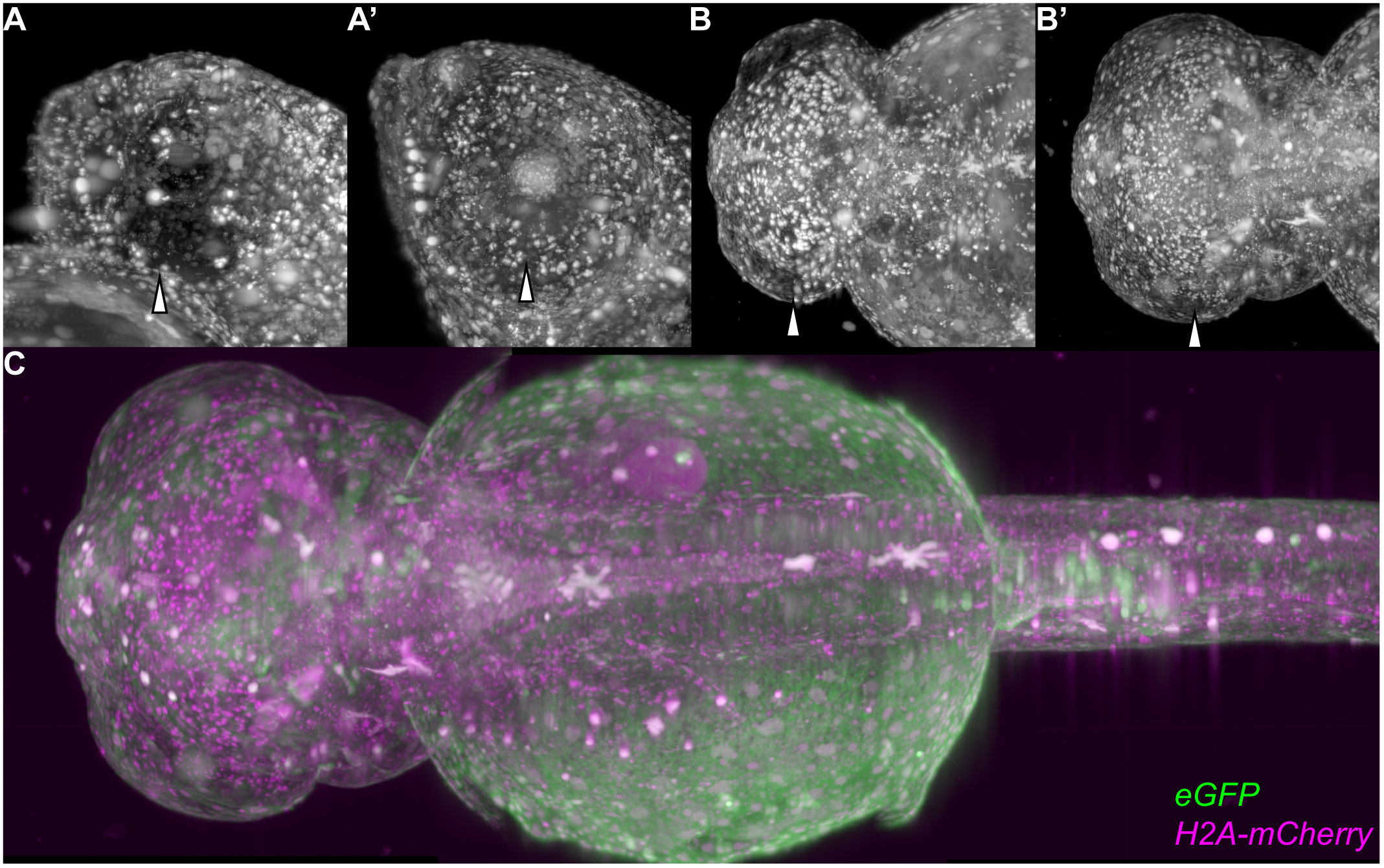
The established toolbox enables imaging of previously challenging organs and tissues, such as the retina and the brain. Using the pigment double knockout in combination with α-Bungarotoxin, eGFP and mCherry coupled to histone2a the imaging was conducted in a single plane illumination microscope (SPIM) we could image the head region of wild-type and *spooky* embryos. Hollow arrowheads point towards the eyes in A and B. (A) Maximum projection of *H2A-mCherry* signal of a lateral view of a wild-type medaka embryo. The eye is not penetrable and no nuclei are visible within or behind it. (A’) Maximum projection of *H2A-mCherry* signal of a lateral view of a *spooky* medaka embryo. The eye is penetrable and nuclei are visible within or behind it. (B) Maximum projection of *H2A-mCherry* signal of a dorsal view of a wild-type medaka embryo. The eye is not penetrable and no nuclei are visible within it. (B’) Maximum projection of *H2A-mCherry* signal of a dorsal view of a *spooky* medaka embryo. The eye is penetrable and nuclei are visible within it. (C) Stitched maximum projections of a dorsal view of a *spooky* embryo. This embryo was imaged for 48 hours, which can be seen in S3 Movie.

## Discussion

In medaka there is a clear trend of fluorescence intensity, which is maintained throughout the imaging time, i.e. ratios between fluorescent proteins remain stable. This allows for the scoring of fluorescent proteins by fluorescence intensity.

The GFP derived fluorescent proteins mVenNB and mGFPmut2 are ranking highest in fluorescence intensity. The most popular and standard fluorescent protein, eGFP, with a considerable toolset available including antibodies, nanobodies, split-GFP, etc. [27–29] only scores number three with respect to fluorescence intensity. Since the eGFP toolset is likely also applicable to mVenNB and mGFPmut2, the balance between necessary tools and absolute fluorescence intensity will determine the final choice. In some instances, optimization and evaluation could be done *in silico* assessing e.g. antibody binding sites or *in vitro* by immunostaining with antibodies or nanobodies. In addition, the establishment of split fluorescent proteins may be possible. Furthermore, both fluorescent proteins carry an A206K mutation, preventing dimerization making those more amenable for fusion proteins [30]. In conclusion, eGFP remains a solid choice if the experiment heavily relies on secondary factors, which are not entirely based on *in vivo* fluorescence readout.

Among the red fluorescent proteins the number one candidate in medaka is mCherry, which is not only the fluorescent protein with the highest fluorescence intensity, but also with the best established toolbox such as nanobodies, antibodies, etc. [31].

Interestingly, when comparing the measured fluorescence intensities to the theoretical values such as quantum yield and excitation efficiency [32], we observed a discrepancy between measured values and expected results. This might be due to the different physiological conditions inherent to each model organism or underlying non-optimal excitation of fluorescent proteins by single wavelengths.

Verification of fluorescent proteins by light-sheet microscopy [1,33] is in line with previous observations. Green fluorescent proteins do differ in contrast to more prominent differences among the red fluorescent proteins. Notably, after mRNA injection fluorescence intensity steadily decreases during development, partially accounting for the small distribution of values. It is important to consider that in transgenic lines or after endogenous tagging by CRISPR/Cas9, the most intense fluorescent proteins are continuously expressed and therefore exhibit their fluorescence intensity and maturation time.

The optimal fluorescent protein in zebrafish is not as easy to score by fluorescence intensity. This is due to varying intensities depending on fluorescent protein and developmental stage of the fish. Here, the choice of the optimal fluorescent protein depends on the experimental condition, including the stage of embryos at the time of analysis. Given that we employed standard cassettes ensuring comparable mRNA stability, this may be due to different kinetics of protein maturation and turnover, affecting the quantitative measurements.

Strikingly, most fluorescent proteins show similar *in vivo* fluorescence after normalization in medaka and *E. coli*. However, some of the tested proteins diverge from this trend, reinforcing the need for testing novel fluorescent proteins specifically in the species of choice. Species specific differences may arise from differences in their translational or maturation machinery, cellular pH-values or any combination of components different between the cells of both species.

In our hands the optimization of codon usages according to establish codon usage tables does not improve, but rather decreases fluorescence intensity [24]. While it had been shown previously in *Ciona intestinalis* that that codon averaging can positively impact on the fluorescence intensity of fluorescent proteins [34]. Our results, in contrast, show that codon-usage table driven averaging is not effective in improving fluorescence intensity, and rather decreased intensity by 25- to 30-fold likely due to changes in mRNA stability or translation efficiency due to repetitive codons.

Even though widely used for fish, the anaesthetic MS222 (Tricaine) has several drawbacks such as non-complete anaesthesia and developmental cardiac defects during extended *in vivo*-imaging [35]. Several groups were therefore trying to supplement Tricaine with different anaesthetics such as Isoflurane [36], Etomidate [37] or increasing the dosage of Tricaine itself [38]. The injection of α-Bungarotoxin as alternative approach has previously been suggested for zebrafish, with apparent influence on development or heart morphology [21]. An obvious concern in long-term imaging experiments is the photostability of the anaesthetics. Since α-Bungarotoxin is delivered as an mRNA, the resulting protein is already at the site of action and will be resynthesized proportional to the amount of initially injected RNA, the number of cell division and the stability of the injected mRNA. The ultimate choice of the anaesthetics to be employed finally depends on the biological process to be studies. If those are affected by α-Bungarotoxin, irreversibly binding to nicotinic acetylcholine receptors [21], Tricaine blocking the sodium channels in neuronal membranes [39] or Etomidate potentiating GABA-receptors in the central nervous system [40] might be valid alternatives with the limitations discussed above.

The pigment double knockout *spooky* is a valuable addition to toolbox since it is readily established in any existing transgenic of mutant background overcoming time consuming complex breeding schemes to re-establish the transparency of e. g. see-through medaka, homozygously mutant for 5 loci. Already the injected generation facilitates instant deep imaging.

The approaches presented here facilitate efficient and deep long-term, *in vivo* imaging of fish embryos and juveniles employing fluorophores with the at best quantum yields in steady, highly transparent embryos e. g. by light-sheet microscopy (Fig 6). We present a streamlined strategy for the optimization of additional species-specific fluorophores and provide cross-validated reference points for immediate comparison of new fluorophores developed.

## Materials and Methods

### Fish husbandry and microinjections

Medaka (*Oryzias latipes*) and zebrafish (*Danio rerio*) stocks were maintained as previously described [41]. All fish are maintained in the closed stocks of COS at Heidelberg University. Fish husbandry and experiments were performed according to local animal welfare standards (Tierschutzgesetz 111, Abs. 1, Nr. 1, Haltungserlaubnis) and in accordance with European Union animal welfare guidelines. The fish facility is under the supervision of the local representative of the animal welfare agency.

Embryos were staged according to Iwamatsu [15]. Medaka microinjections were performed as previously described [3].

### CRISPR/Cas9 and single guide RNAs

JDS246 was a gift from Keith Joung (Addgene plasmid # 43861). Cas9 mRNA was transcribed by mMessage mMachine Sp6 Transcription Kit (Thermo Fisher). Single guide RNAs were predicted by CCTOP [24] and subsequently cloned into DR274 [42]: sgRNA(Oca2_1): GAAACCCAGGTGGCCATTGC, sgRNA(Oca2_2): TTGCAGGAATCATTCTGTGT, sgRNA(Pnp4a): CCTGGGGGCGGTGCTTAGTC. DR274 was a gift from Keith Joung (Addgene plasmid # 42250) [42]. The vector was linearized by DraI-digest and transcribed by MEGAscript T7 transcription Kit (Thermo Fisher). All RNAs were purified by RNeasy Mini Kit (Qiagen). Injected concentrations were: 150 ng/μl Cas9 mRNA and 15 ng/μl per sgRNA.

### Loading and imaging of 96-well plates

96-well plates were loaded with 150 μl volume per well, including the treatment solution and the embryo. The plates were loaded based on a randomized loading scheme created by a customized R script, which creates a loading scheme and a table of each well for further use in the analysis. The plates were sealed by a gas permeable moisture barrier seal (4ti-0516/96, 4titude, Dorking, England), loaded into the ACQUIFER Imaging Machine (DITABIS, Pforzheim, Germany) and imaged for a defined time and channel composition. The plates were continuously incubated at 28°C. An overview of the imaged plates can be found in S2 Table along with the timeframe, timestep and used channels. A general description of the imaging condition can be found in the section about the semi-automated analysis of the experiment, i.e. either anaesthetic or fluorescent protein analysis.

The used LEDs and filters in the ACQUIFER Imaging Machine were a 470 nm LED in combination with LED-FITC-A-000 (Semrock, Inc.) for the green channel and a 550 nm in combination with a LED-TRITC-A-000 (Semrock, Inc.) for the red channel.

### Anesthesia reagents

α-Bungarotoxin mRNA was transcribed from pmtb-t7-alpha-bungarotoxin, which was a gift from Sean Megason (Addgene plasmid # 69542) [43]. The vector was digested by EcoRV-digest and the linearized plasmid transcribed by mMessage mMachine Sp6 Transcription Kit (Thermo Fisher) for 4h. The mRNA was purified with the RNeasy Mini Kit (Qiagen). Embryos were microinjected with 3-25 ng/μl α-Bungarotoxin mRNA along with 50 ng/μl eGFP mRNA. Embryos were then either mounted into a 96-well plate for assaying anaesthesia efficiency or the number of hatched and swimming embryos was scored for developmental stages.

Etomidate solution was prepared as a 10 mg/l solution (7.5 μl of a 4 mg/ml stock solution in DMSO in 3 ml 1x ERM). The DMSO solution was prepared accordingly, yielding a 0.25% DMSO solution. A solution of 1x Tricaine (0.02% w/v, A5040, Sigma-Aldrich) was prepared.

### Heart morphology scoring and heart rate measurements

Embryos were imaged for 10 seconds with 25 frames per second in order to score heart morphology and heart rates at 25°C. The data was imported into R [23] and plotted using ggpubr and xlsx libraries [44].

### Startle-response assay

Demounted, with α-Bungarotoxin mRNA injected embryos and control embryos were startled with a pipette tip for 10 times. The number of responses was noted and plotted in R using the ggpubr library.

### Fluorescent protein cloning and transcription

All fluorescent proteins were cloned into pGGEV3 from the Golden GateWAY toolkit [45]. Each construct was cloned with a start and stop codon to ensure maximum comparability of the resulting mRNAs. Fluorescent proteins were cloned from present plasmids except for the following. cytoplasmic EKAR (Cerulean-Venus) was a gift from Karel Svoboda (Addgene plasmid # 18679) [46], mRuby2-C1 was a gift from Michael Davidson (Addgene plasmid # 54768) [47], pcDNA3-Clover was a gift from Michael Lin (Addgene plasmid # 40259) [47], pmScarlet-i_C1 was a gift from Dorus Gadella (Addgene plasmid # 85044) [43] and SceGFP was a gift from Sabine Strahl [48] eGFP was codon optimized for medaka by OPTIMIZER [19] resulting in the sequence of OleGFP. OleGFP was synthesized by GeneArt Gene Synthesis (Thermo Fisher). Cloned plasmids were linearized with SpeI prior to transcription and transcribed by mMessage mMachine Sp6 Transcription Kit (Thermo Fisher). Fluorescent protein mRNA was microinjected in equimolar amounts compared to 50 ng/μl eGFP mRNA.

### SPIM image acquisition

Medaka embryos were anesthetized with 1x Tricaine or by α-Bungarotoxin mRNA injection and stacks were acquired in a MuVi-SPIM (multiview selective plane illumination microscope) [13,14,49] as described before [50] for the 25x detection setup, with the addition of a 488 nm illumination and a corresponding 525/50 bandpass filter. For the measurements of fluorescence intensity the resulting images were divided by the control channel, masked by the Otsu algorithm [51] and subsequently measured.

The resulting tables were imported into R [23] and fused into one dataset. The measurements were linked to conditions and plotted using ggplot2 [52]. The used R libraries were: data.table [53], zoo [54], ggrepel [55], and ggpubr.

The stacks resulting in Fig 6 were recorded and imported into Fiji [22]. They were transformed with the logarithm due to the varying expression of cell types and autofluorescence of leftover pigments.

### Comparison of medaka and *E. coli in vivo* fluorescence intensity

*E. coli* values were extracted from supplementary material of Balleza et al. [16]. Medaka values were calculated as a mean of all data of one fluorescent protein from 23 to 30 hpf from imaged data and normalized to mRFP1* for red fluorescent proteins, eGFP for green fluorescent proteins and Venus for yellow fluorescent proteins (set to a value of 1.2, according to the *E. coli* value).

### Codon usage table driven codon averaging and adaptation index

Codon averaging was performed by OPTIMIZER [19], using the codon usage table, which can be obtained from HIVE-CUTs [56]. Codon adaptation indices of all used sequences were scored by using the seqinr library in R [57] and codon usage tables from HIVE-CUTs [56]. The data was plotted using ggplot2 [52], readr [58], data.table [53], condformat [59] and ggrepel [55].

### Semi-automated analysis anesthesia

One slice of each well was acquired by autofocus (16 steps with a distance of 25 μm, software-based) per timepoint in brightfield with a CFI Plan Achromat UW 2x, N.A. 0.06 objective (Nikon). Illumination was performed for 20 ms at an LED intensity of 100 %. The timelapse was recorded for at least 45 hours in 20 or 33 minute intervals. Subsequently the imaged medaka were staged for both uösed plates and the timepoint 0 was set according to the higher starting stage (stage 28) in order to be able to fuse the plate data. The following steps were performed in Fiji [22]. These images were concatenated to a stack per well. Subsequently the square of the difference of timepoints was calculated and exported as a table.

The resulting tables were loaded into R [23] and fused into one dataset. The negative wells (dead or non-developing embryos) were dropped from the data set, the wells were linked to the corresponding conditions and the data was plotted using ggplot2. The used R libraries were: data.table [53] and ggplot2 [52]. Customized scripts are available through github (https://git.io/fAPnh).

### Semi-automated analysis of fluorescent proteins

A total of 12 z-slices with 100 μm step size were acquired with a CFI Plan Fluor 4x, N.A. 0.13 objective (Nikon) per well, guaranteeing the embryo to be included in the image. These slices were acquired in the previously described green and red channel. Illumination was performed for 50 ms at an LED intensity of 50 % for both channels. Yellow and green fluorescent proteins were recorded in the green channel due to the limitation of light-sheet microscopes of requiring the according filter. The timelapse was recorded from directly after injection up to at least 40 hours for the longer experiments, once each day for the 2, 3 and 4 dpf experiment and from injection to 48 hours for the codon adaptation experiment. The following steps were performed in Fiji [22]. Each slice of each channel was masked by both channels by applying an auto threshold with the Otsu algorithm and cropping each channel by both resulting masks [51]. The resulting, masked images were measured by excluding the background. All wells were checked for fluorescence at the maximum and the half-maximum timepoint in order to exclude developmentally impaired or dead embryos.

The resulting tables were imported into R [23] and fused into one dataset. The negative wells were discarded, the wells were linked to the corresponding conditions and normalized to the injection control per well at 10 hpf. Used was the data masked by the channel of the injection control to ensure automatic cropping. The data was plotted using ggplot2. The used R libraries were: data.table [53], zoo [54], ggrepel [55] and ggplot2 [52]. Customized scripts are available through github (https://git.io/fAPnh).

## Acknowledgements

We thank Jochen Gehrig and ACQUIFER (DITABIS AG, Pforzheim) for cooperation on automated microscopy and technical feedback. We thank Alex Cornean, Clara Becker, Tinatini Tavhelidse, Jakob Gierten and Thomas Thumberger for critical reading and comments on the manuscript. We also thank Dimitri Kromm and Lars Hufnagel for continuous support in light-sheet imaging. We thank Alex Cornean for contributing to the schemes in Fig 1 and 4. We thank Steffen Lemke, Lars Hufnagel and Thomas Höfer for critical comments on the manuscript.

## Data availability

Not available, raw data used exceeds the reasonable limits of Dryad and Figshare (> 11 TB).

## Supporting information

**S1 Fig. Wild-type vs. dechorionated fluorescence extraction.** Embryos were injected as previously mentioned. Half of each condition was dechorionated. Both conditions were imaged at 2, 3 and 4dpf and compared to each other. Optical properties of the chorion were negligible and therefore neglected in the analysis since dechorionation has a larger impact on development, especially when performed in an early stage (2-4 dpf).

**S2 Fig. Codon adaptation indices of all used sequences.** Codon adaptation index of each sequence is plotted for medaka and human along with a diagonal line for orientation.

**S3 Fig. Single well projections along time.** Single wells were projected with three different projection types along time: standard deviation, average projection. In case the embryos are not moving it is to be expected to see the full contours of the embryo in the standard deviation and in the average projection. This is not the case for most of the conditions, except for the α-Bungarotoxin mRNA injected embryo.

**S4 Fig. Injections of α-Bungarotoxin have no detectable impact on heart development.** (A) Embryos after demounting from the 96-well plate and finishing the assay (stage 31). No cardiac defects are visible in both controls and embryos injected with α-Bungarotoxin mRNA. For Tricaine, mild defects can be detected, whereas severe defects can be observed in Etomidate treated embryos. (B) Heart rate of α-Bungarotoxin is statistically indistinguishable to its control (mock injection). Mild defects and changes can be observed in Tricaine and Etomidate treated embryos (asterisks indicate P-values: **** P <= 0.0001, *** P <= 0.001, ** P <= 0.01, * P <= 0.05, ns P > 0.05). (C) Different dilutions of α-Bungarotoxin were tested in order to adjust the concentration so fish would wake up after microscopy. Embryos were injected with 0, 3, 6, 12 or 25 ng/μl α-Bungarotoxin mRNA along with eGFP mRNA as an injection control. The number of hatched embryos and the number of swimming embryos per condition was scored (0 ng/μl: n = 33 fish, 3 ng/μl: n = 34 fish, 6 ng/μl: n = 16 fish, 12 ng/μl: n = 24 fish, 25 ng/μl: n = 22 fish).

**S5 Fig. Adult medaka with pigment knockout.** Addition to Fig 5B. Stage 27, hatched and juvenile fish of the wild-type strain and the double pigment knockout line were imaged. In addition, *oca2^-/-^* fish are shown at stage 36 and adult.

**S1 Movie. Embryos with different anesthetics.** Collection of embryos used in the analysis of Fig 4B. Color code is matching the color code in Fig 4.

**S2 Movie. Development of a wild-type and mutant embryo.** Development of a wild-type and a *oca2^-/-^* pigment knockout embryo.

**S3 Movie. A developing spooky embryo.** Stitched stacks of a developing *spooky* embryo, injected with with *α-Bungarotoxin*, *eGFP* and *mCherry* coupled to *histone2a.* The imaging time was 48 hours and the time interval 1 hour between the stacks.

**S1 Table. Fluorescent proteins used in this study.** All fluorescent proteins used throughout this work including addgene identifiers for comparisons, sequences and mutations from ancestor fluorescent proteins. Furthermore, fluorescent proteins were compared to the classification in the recent publication of Balleza et al., where they made an effort to distinguish the different fluorescent protein versions [16].

**S2 Table. All plates analyzed for this study.**

